# A Combination of Environmental and Landscape Variables Drive Movement and Habitat use in Two *Anaxyrus* sp. in the Eastern Coastal Plain

**DOI:** 10.1101/2024.02.29.582809

**Authors:** Alexander M. Ferentinos, Courtney E. Check, Olivia Windorf, Matthias Leu

## Abstract

Amphibians are one of the most endangered taxa and are largely threatened by habitat loss. Little work has been conducted on the movement and habitat use of amphibians outside of the breeding season. In this study, we examined the movement patterns of two species of toads inhabiting the Eastern Coastal Plain of Virginia: the Eastern American Toad (*Anaxyrus americanus americanus*) and the Fowler’s Toad (*Anaxyrus fowleri*). Based on three years of movement data, we estimated the median migration distance of toads from their breeding location and the propensity for site fidelity, related variation in distance traveled to environmental (e.g., rain, temperature, humidity) and landscape variables (e.g., coniferous forests, distance to trails, terrain ruggedness index), and compared microhabitat selection for daytime refugia between the two species. We found the median distance from breeding grounds to be similar between the two species, 63 m for the Eastern American Toad and 64 m for the Fowler’s Toad, but Eastern American Toads had a greater range of moved distances (3^rd^ quartiles were 122 m for Eastern American and 73 m for Fowler’s Toads). We also found that both species exhibit site fidelity. Distance to trails, proportion of conifer forest, minimum temperature, and 3-day cumulative rainfall related positively with increased movements. Compared to Fowler’s Toads, Eastern American Toads favored woody structures and leaf litter for daytime refugia. Our research provides crucial information for two toad species about the extent of their movements and habitat use during the nonbreeding season. To lessen the decline of amphibians, habitat occupied during the nonbreeding season needs to be included in conservation strategies at biologically relevant distances.

---

Habitat loss is one of the most prevalent drivers of extinction across all taxa (Leu et al., 2019), and in amphibians, the removal of habitat, such as forests and wetlands, reduces connectivity and population recruitment. Connectivity and dispersal are essential for the continuation of a species in an area to combat periodic local extinctions and to maintain genetic diversity (Cushman, 2006). Habitat loss can also skew sex ratios due to differential dispersal patterns between sexes outside of the breeding season. This means loss of habitat at certain extents can severely reduce the presence of one sex in the area, thereby limiting the reproductive success, or resulting in overcrowding of now-limited habitat (Johnson et al., 2007).

Amphibians are one of the clades of vertebrates most suffering the greatest declines in from the Anthropocene, as 41% of assessed amphibians are currently at risk of extinction (International Union for Conservation of Nature, 2023). The vulnerability of amphibians to habitat loss is especially notable given their biphasic lifestyle. Amphibians frequently rely on shallow ponds as breeding sites which exclude aquatic predators, such as fish, but allow larvae access to ample food supplies (Hecnar & M’Closkey, 1997; Knutson et al., 2004). Amphibian migrations to breeding sites vary greatly among and within species, with adult terrestrial species migrating mean distances of 142 - 289 m to reach breeding sites. While adult movements are relatively stable, both dispersal to new sites and local population stability is hypothesized to be supported by juvenile movement from breeding sites after metamorphosis (Semlitssemch, 2008). Therefore, adult abundance acts as an indicator of non-breeding habitat quality, as amphibians have been found to occupy sites based on critical thresholds in the amounts of their preferred habitat (Regosin et al., 2005; Trenham & Shaffer, 2005). However, amphibian movement and habitat use during the non-breeding season are understudied (Bailey & Muths, 2019).

The loss of amphibian diversity is a concern for the world at large. Amphibians broadly occupy a key middle trophic role, where they both control pests that exist at lower trophic levels, such as mosquitos, and act as a food source themselves for a variety of more visible and monetizable species, such as birds and fish. At an ecological level, amphibians act as key bio-indicators of water and habitat quality due to the permeability of their skin (Cortés-Gomez et al., 2015; Pollet & Bendell-Young, 2000). Finally, amphibians are of sizeable interest to humans, both as a culinary item and for medical research. They possess numerous abilities, such as the ability to regrow tails and to turn off stomach acid production, which make them important models in the study of regeneration and digestive issues (Hocking & Babbitt, 2014). Therefore, understanding the movement patterns and habitat use of amphibians during the non-breeding season will aid in their conservation thereby ensuring their contribution to ecosystem functioning and services (Pittman et al., 2014).

Here, we study the movement patterns and habitat use during the non-breeding season of two representative species of upland utilizing anurans in the Coastal Plains of Virginia: the Eastern American Toad and Fowler’s Toad. Eastern American Toads are a widespread species found throughout northeastern North America (IUCN SSC Amphibian Specialist Group, 2020), while Fowler’s Toads occupy most of the eastern half of the United States (US hereafter; IUCN SSC Amphibian Specialist Group, 2021). While neither is considered globally or federally endangered, Fowler’s Toads are a species in decline in localized areas, as populations across the state of Virginia have fallen by approximately 53% (Jones & Tupper, 2015), and in Ontario, Canada, and several US states at the periphery of its range, it is considered locally threatened (Vermont Fish & Wildlife Department, 2022; Yagi & Green, 2017). Additionally, the human population in our study area in southeastern Virginia has increased over the last several decades, exerting more stress on local populations (U.S. Census Bureau, n.d.). Protections to amphibians in Virginia are mainly provided by the requirement of an approximately 30 m (100 ft) buffer around Resource Protection Areas (Virginia Department of Wildlife Resources, 2021).

This study used a combination of harmonic direction-finding (HDF) and radiotelemetry relocation techniques to examine the environmental and landscape level drivers of movement, the distance migrated from breeding sites, microhabitat use, and site fidelity in Eastern American Toads and Fowler’s Toads between late spring and late summer. Previous studies have provided a variety of migration distances in American Toads (Bartelt & Klaver, 2017; Forester et al., 2006; Oldham, 1966), but less literature exists on Fowler’s Toads, with work generally limited to an isolated population in Ontario (Boenke, 2012; Smith & Green, 2006). We focus on these species’ movements and habitat use in the coastal plain of the eastern seaboard, where neither species has been intensively studied before. This research will allow conservation managers to establish whether the space allocated to amphibian populations in human landscapes such as parks, ponds and wetlands with vegetative buffers is adequate to their needs (Semlitsch & Bodie, 2003).

## MATERIALS AND METHODS

### Study areas

Our study area consisted of 4 separate locations in Williamsburg, VA. The first was the College Woods of the College of William & Mary (37.2752, −76.7234), composed of primarily mixed beech (*Fagus* spp.) and maple (*Acer* spp.) forest with several shallow wetlands surrounding a large reservoir lake, and includes several graveled walking trails. The Greensprings Interpretive Trail site (37.2499, −76.7913; hereafter referred to as Greensprings Trail), consists of mixed hardwood and conifer forests surrounding a beaver-maintained wetland, and includes several well-maintained walking and biking trails. The Greensprings Field site (37.2397, −76.7868) is located on a portion of the Virginia Capital Bike Trail to the southeast of the Greensprings Interpretive Trail Site and was only included in the study in 2023. This site was bordered by a forest and by the Historic Mainland Farm, which mainly grows corn, wheat, and soybeans. The ground on both sides of the path has depressions and ditches which fill with water during rain events and are used as breeding sites by amphibians. The Warhill Sports Complex site (37.324, −76.7603; hereafter referred to as Warhill), consists of a large reservoir, surrounded by mixed deciduous-conifer forest, along with a well-maintained set of hiking trails.

#### Construction & Attachment of HDF Transponders and radio-telemetry belts

Both kinds of belts were constructed based on a design previously tested by Alford & Rowley (2007). HDF transponder belts were constructed from medical grade silicon tubing (1 mm inner diameter x 2 mm outer diameter) of approximately 10-15 cm in length to accomodate larger individuals. We used medical tubing to increase the surface area of the force exerted by the binding cotton thread, preventing abrasions from developing on the skin of the toad due to friction, which are the most frequent injury from external radio telemetry transponder attachment in amphibians (Groff et al., 2015). We tested this method with Eastern American Toads and Fowler’s Toads in a pilot study (Check, 2019; Windorf, 2019). It also allowed for a natural release mechanism of the belt due to degradation of the cotton thread. At the middle point of each belt, we used a push pin and X-ACTO knife (Elmer’s Products Inc., Westerville, OH) to create a slit for the transponder (RECCO, Lidingö, Sweden) to be inserted into with the central diode hanging out on one side to prevent coverage of the diode, which could potentially weaken the signal. A dab of Aquarium Safe All-Purpose Adhesive Sealant (DAP Product, Inc., Baltimore, MD) was applied to the top side of the tag, with care being taken to not cover the diode. To identify individual toads once detected, we used unique color-coding schemes painted on the silicon belts with enamel gloss acrylic paints (FolkArt, Norcross, GA), which we found to be resistant to water and outdoor conditions in a pilot study (Check, 2019; Windorf, 2019). Color codes were generated with SalaMarker (MacNeil et al., 2011), for permutations of Visible Implantable Elastomers. This code was used to generate paint permutations, with 8 paint colors and 2 locations (left and right of the transponder, with the diode facing the viewer) available, generating 64 unique codes.

Radio transmitters (Advanced Telemetry Systems, model 1025, 0.65 g, battery life of 45 days) were attached via a similar method to the HDF transponders. We constructed harnesses for toads in the field to maximize battery life. When a toad of appropriate weight was captured, a length of tubing was fitted around its waist while the toad was restrained. This length was then cut off, and a section in the middle of the belt that was the width of the transmitter was cut out. We then inserted a cotton thread through all 3 components, creating a secure belt that was tied around the toad with square knots. Trimming of the silicon tubing to ensure a snug fit occurred as with the HDF belts. We only affixed tracking devices to individuals whose body weights were at least 20 times larger than the weight of the device, in keeping with the 5% body weight rule (Goldberg et al., 2002; Z. T. Long et al., 2010).

To capture toads, pairs of researchers went out at night following a rain event, when the toads were most active, and walked paths where toads were known to be, capturing those we came across (Clark, 1974; Fitzgerald & Bider, 1974). For each capture, we georeferenced location with a GPS unit (Garmin GPSMAP 64s Olathe, KS)), and noted capture substrate, specimen weight using a plastic bag and a 50 g spring scale (Pesola, Schindellegi, Switzerland), and snout-vent length (SVL, mm). Capturing and tagging toads was approved by the William & Mary’s IACUC (permits 2023-0095, and Protocol IACUC-2017-02-20-11745-mleu) and the Virginia’s Department of Wildlife Resources (permits 059643 and 3581301).

### Relocation Methods

To relocate individuals every 1 to 3 days (average number of days between relocation events), we started at the last known location of a tagged individual. For the HDF methods, we turned on the RECCO R9 Detector (RECCO, Lidingö, Sweden) to detect the specimen on the approach. Upon reaching the last known location, we scanned the immediate surroundings for a signal. If no signal was received, we engaged in a serpentine search pattern of 4 quadrants that were at maximum 10 m x 10 m. Within each quadrant we walked 2-3 m wide transects, as the potential range of the detector was hampered by leaf litter, coarse woody debris, trees, or other obstacles.

Once a signal was received, we homed in on it by turning down the volume of the detector. We determined the exact location by carefully sifting through obscuring debris to get visual confirmation of individuals’ color combination. We recorded recapture location with Garmin GPSMAP 64s, refugia substrate, and time of day. To reduce stress on toads, visual confirmation was only considered necessary for recaptures in new areas, or after 1 week of no movement, as previous ones were assumed to be the specimen remaining in its refugia. If a given individual was not relocated, it was marked as a No Capture (NC) for that day, and if one full week of NC occurred, it was marked as lost. We also considered an individual lost if only the belt and transponder were found, and recorded all other information as if it was a recapture.

#### Combined Radiotelemetry and HDF Technique

While typically presented in opposition to one another (Alford & Rowley, 2007), in this study, we found the combination of radiotelemetry and HDF techniques result in more efficient relocation times. This is because radiotelemetry and HDF are complementary to one another in this study. Radiotelemetry works well at long range, but is imprecise at short ranges, especially when working with small animals that have cryptic behaviors. HDF on the other hand, has limited long range capabilities, as material between the receiver and the diode on the tag can “muffle” the signal, and limit its advertised range.

In combining them, we first began by going through the typical methods of radiotelemetry relocation, approaching areas with the highest signal strength, before turning down the gain, and repeating until they arrived at a search area (Fuller & Fuller, 2012). While simple triangulation could give an idea of what area to search, it did not provide the exact location of the transmitter. By utilizing the RECCO HDF detector, we could pick up a signal from the radio telemetry transmitter on toads’ backs and acquire their precise location. This was possible because while HDF tags are easiest to detect with the RECCO detector, we found that many electronic/metal objects also returned a signal and could be located.

### Covariates used in the analyses

We evaluated variation in distance moved on the basis of time of year, spatial and weather data. We included time of year since previous research has shown that frog movement varies in relation to time of year (Clark, 1974; Groff et al., 2017), and because the movement data were exclusively post breeding for the Eastern American Toad (Breeding season in the Carolinas: February - March), but included movement during the breeding season for the Fowler’s toad (Breeding season in Virginia: April – July; Virginia Herpetological Society, 2023). We included spatial data such as terrain ruggedness because previous research showed that rugged terrain was less favorable for movement to amphibians (Homola et al., 2019; Urban et al., 2008). We included land cover data, such as deciduous and coniferous forest cover, because studies have shown that Eastern American Toads frequently make use of woody debris in forests for shelter, while Fowler’s Toads generally favor more open habitat types (Jones & Tupper, 2015; Pitt et al., 2013; Semlitsch et al., 2008). We included anthropogenic features, such as distance to trail, because we observed that toads were frequently on trails at night, and González-Bernal et al. (2011) found that toads made more attempts to capture prey on path substrates.

Daily temperature rainfall and relative humidity were included because our pilot study (Check, 2019; Windorf, 2019) suggested them to be important and because previous research showed that increases in these variables were associated with greater movements by toads (Greenberg & Tanner, 2005; Phillips et al., 2007; Sinsch, 1988). We included distance from breeding grounds and water bodies because we were interested if there was any effect on proximity to them to the movements of toads, even outside of the breeding season.

#### Spatial data

We developed a terrain ruggedness index from the national Digital Elevation Model data at the 1-m resolution (U.S. Geological Survey, 2023a). To create a Terrain Ruggedness Index (Riley et al., 1999) raster, we used the “Ruggedness Index” tool in QGIS (QGIS Development Team, 2023).

Land cover data were obtained from the Virginia Land Cover Dataset, available at 1-m resolution (Virginia Geographical Information Network, 2023). We reclassified land cover data by combining fine-resolution categories, such as levels of imperviousness, into coarse-level categories (for crosswalk see Supplemental Table 1). Using images from NASA Planet Labs at 3-m resolution, we classified coniferous land cover on the basis of winter scenes (Marta, 2018), using the Supervised Classification tool in ENVI version 5.7 (Exelis Visual Information Solutions, 2023). To match resolutions, we resampled The Virginia Land Cover Dataset to a 3-m resolution using the “Nearest” method. We combined the Virginia Land Cover Dataset and coniferous land cover dataset using the “Raster Calculator” tool.

To calculate distance from trails, water, and breeding sites, we used the “Euclidean Distance” function in ArcGIS Pro version 3.2 (ESRI, 2023). Unless stated otherwise, this was the program used for spatial analyses. Shapefiles for trails in the College Woods (McKinney, 2002), Warhill (*Warhill Trail*, 2023), and the southeastern loop of the Greensprings Trail were available online (*Greensprings Interpretive Trail*, 2023). We hand-digitized the trail for Greensprings Field on the basis of aerial imagery, and delineated the northwestern loop of the Greensprings Trail in the field using the Track function on the Garmin GPSMAP 64s. Delineations of bodies of open water were taken from a combined dataset of both open water based on the Virginia Land Cover Dataset, as well as small streams based on the National Hydrography Dataset (U.S. Geological Survey, 2023b). We hand digitized several streams and water bodies that were absent from these datasets.

These land cover data, including the proportion of forest and coniferous forest, were sampled with moving windows at the 1^st^ quartile, median, and 3^rd^ quartiles of non-zero movements by all toads. Exponential decay functions were also fitted to the distances of toads from trails, water, and breeding sites to investigate non-linear relationships.

#### Weather data

Daily maximum, mean, and minimum temperatures, rainfall, and mean dew point were obtained from the PRISM data set, which is available at the 4-km resolution (Oregon State University, 2023). Relative humidity was calculated on the basis of the mean temperature and mean dewpoint temperature (National Weather Service, n.d.), and rainfall was transformed into cumulative totals ranging from 1-7 and 14 days, as a pilot study found that cumulative totals were more informative than daily ones (Check, 2019; Windorf, 2019). Weather data were joined to dates with movement with a lag of 1 day, as toads primarily moved at night, indicating that the previous day’s weather was a better predictor of movement (Clark, 1974; Fitzgerald & Bider, 1974).

### Data Analysis

#### Distance moved

The distances between subsequent relocations of toads were calculated in R, version 4.3.1 (R Core Team, 2023) using the package “sf” (Pebesma, 2018). All statistical analyses were conducted in R. We did not include the capture location and first relocation for each toad in this analysis to avoid increasing variation in distances moved due to nighttime activity and the impacts of potential fear-driven movements due to tagging. We then used negative binomial generalized linear mixed-effects models from the “lme4” package in R (Bates et al., 2009) to relate variation in distances moved to species (binary variable with Eastern American Toad set to 1), land cover, distance, and environmental covariates. Site and year were included as a random intercept as we expected distance moved to differ among sites and years. We also included Julian day as we predicted that movement would vary over the summer. Each land cover covariate was independently examined across three extents (1^st^ quartile [3 m], median [7 m] and 3^rd^ quartile [17 m] of daily movements), linear vs. decay distance, and against the null model (i.e., intercept model). We used Akaike Information Criterion (Burnham & Anderson, 1998) to select candidate variables, and removed a candidate variable if the null model was the top model or was within 2 AIC units of the top model. All combinations of top candidate variables, including species and Julian day, were evaluated with AIC with the dredge function of the “MuMin” package and effect sizes and SEs were model averaged to derive a final model (Bartón, 2023). The residuals for the final model were evaluated with the “DHARMa” package (Hartig & Hartig, 2017).

#### Site fidelity

We used the “fidelity” package in R (Picardi et al., 2023) to estimate site fidelity on the basis of the Blischke-Scheuer method-of-moments estimator, which calculates the step length parameters of shape and scale for a comparison with simulated toad movements according to a Correlated Random Walk (CRW). We included every toad with 3 or more relocations after the removal of the initial capture points in this analysis. The Blischke-Scheuer method-of-moments estimator was used due to the presence of zeros in the movement between observation data (Rinne, 2008). Movement distances between observations were used instead of the averaged daily distance because the “fidelity” package relies on observation number and order, not dates to estimate the distance a simulated toad should move. Utilizing averaged distances would produce underestimates of the movement ranges in the simulated toads. We defined the distance threshold for considering a movement to be a return to a previous site as 3.65 m, which was based on the margin of error of the Garmin GPSMAP 64s (GARMIN, 2023). A toad had to be absent from a previously occupied site for one observation day before returning to be considered a return. We compared the number of returns, standardized by the number of steps of each toad, first between the observed and simulated movements within the species derived from CRW with paired t-tests. We then compared the observed movements of the two species with a t-test. We set the level of significance to p = 0.05.

#### Substrate analysis

We compared substrate use between species with generalized linear mixed-effects models in the “lme4” package, with site set as a random intercept. The dependent variable was a binary species variable with Eastern American Toad set as 1, and the independent variable was the substrate type. To reduce the impact of capture responses on daytime refugia, we used the same points as in the distance moved analysis. Substrate variables were collated into the major substrate categories (see Supplemental Table 2 for crosswalk). We used the same statistical approach to estimate the final model as outlined for the final distance-moved model.

#### Comparison of radiotelemetry and HDF

We evaluated the differences between radiotelemetry and HDF with three different metrics: 1) the distance between the initial capture and first relocation point, 2) the maximum detected movement between relocations, and 3) the number of days tracked. We used generalized linear mixed-effects model in the “lme4” package with site set as the random intercept. The independent variables in the first two analyses were tag type, species, and weight, whereas in the third analysis we only examined tag type. We report effect sizes ± 95% CI, and point estimates as median (1^st^ and 3^rd^ quartile) unless noted otherwise.

## RESULTS

Sample size for the movement and substrate analyses was 632 for Eastern American Toads and 865 for Fowler’s Toads. The site fidelity analysis was based on 36 Eastern American Toads and 48 Fowler’s Toads.

### Distance traveled from breeding sites

The median distance of Eastern American Toads from their breeding sites was 63 m (1^st^ quartile – 3^rd^ quartile: 43 m - 122 m). The median distance of Fowler’s Toads from their breeding sites was 64 m (1^st^ quartile – 3^rd^ quartile: 31 m – 73 m).

### Movement analysis

We examined 13 candidate variables as drivers of movement (Supplemental Tables 3-4), but included only six in our final analysis, with the remaining seven variables having higher AIC values than or were within 2 ΔAIC from the null model. The top predictors of movement, sorted from most to least important were the minimum temperature, distance to trails, the cumulative three-day rainfall on the day prior to movement, and the proportion of coniferous at the 17-m extent. In regard to species, the negative slope indicates that Fowler’s Toads respond more strongly to the aforementioned variables than Eastern American Toads (Fig. 2). The residual analysis of the final model had a dispersion value of 0.54 with a p-value = 0.008. However, based on the low dispersion value, along with the appearance of the QQ plot and the residual vs predicted graph (see Supplemental Figure 1), we concluded this to be a good model. We believe the significant dispersion value is due to our large sample size.

**Figure 1.**
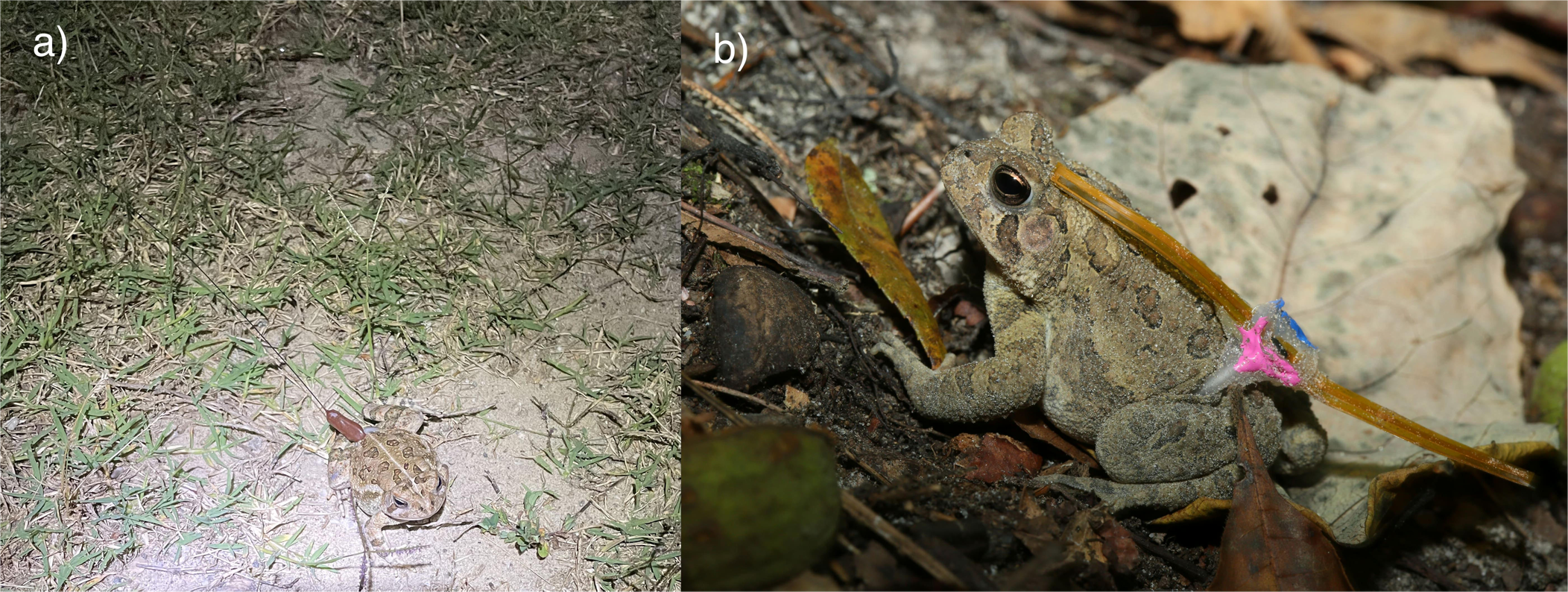
a) Radiotelemetry belt and b) HDF attached to an *Anaxyrus fowleri*. Silicon tubing that contains cotton thread is visible on both toad’s left side. a) taken by Alexander Ferentinos, b) taken by Stephen Salpukas.

**Figure 2.**
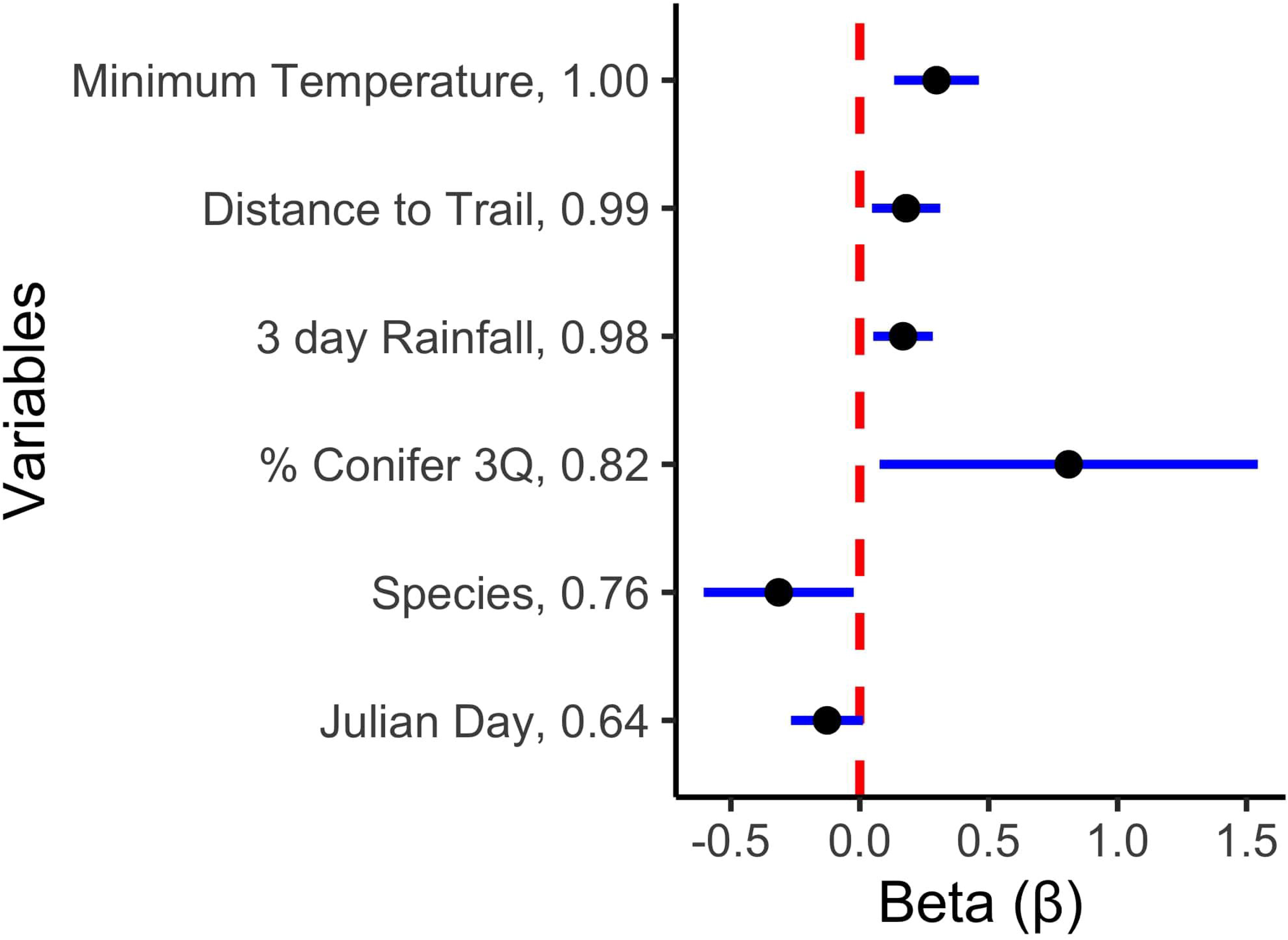
Model-averaged effect sizes and 95% confidence interval for all variables included in the final model of toad movement. Variables are sorted from greatest to least AIC_c_ weight, which is reported after variable name. For species, *Anaxyrus americanus americanus* was set to 1.

### Site fidelity

Both Eastern American Toads (t = 3.04, p = 0.004) and Fowler’s Toads (t = 3.98, p=0.0002) exhibited site fidelity compared to CRW simulated site fidelity data (Fig. 3). Site fidelity did not differ between the species (t = 0.67, p = 0.51).

**Figure 3.**
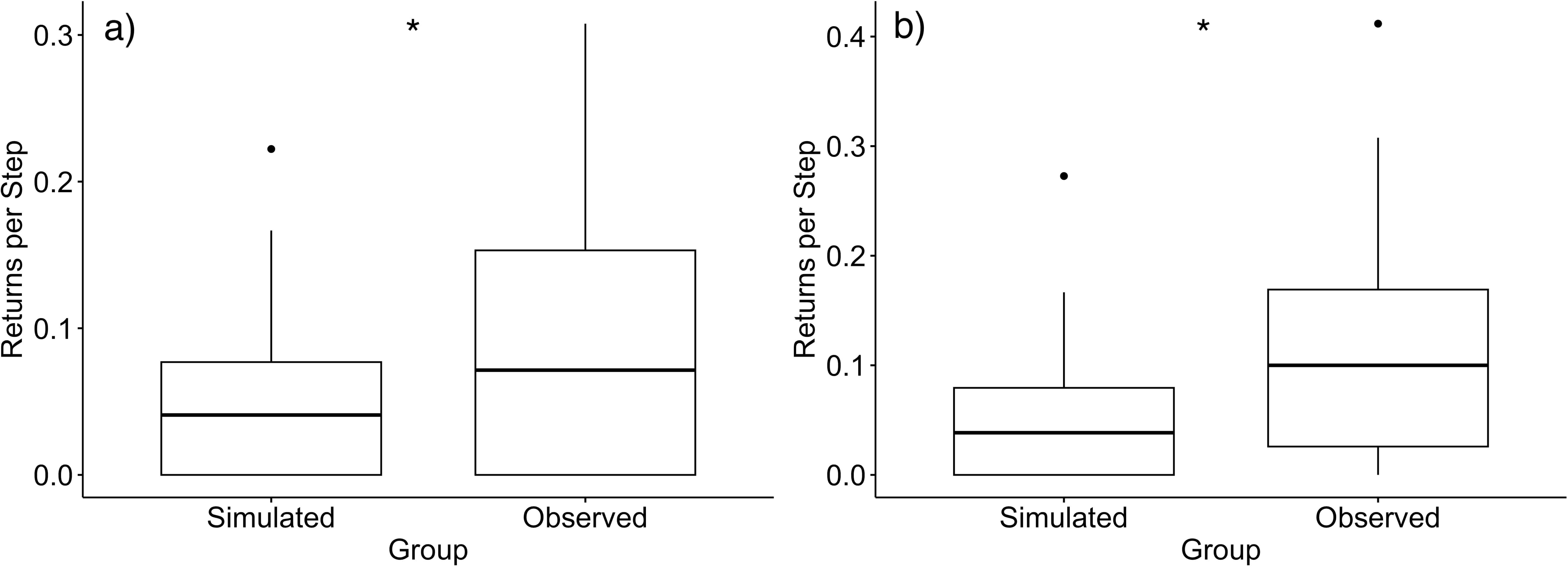
Boxplots of a) *Anaxyrus americanus americanus* and b) *Anaxyrus fowleri* site fidelity as measured by returns per observations in simulated (null) versus observed (obs) toad movements. Horizontal lines represent 1^st^ quartile, median, and 3^rd^ quartiles. Asterisks indicate significance (p<0.05).

### Substrate analysis

We began with 8 variables related to refugia substrate, and after testing them individually, were left with 4 for our final model (Supplemental Table 5). After model averaging, our analysis of substrate choice indicated that the top predictors of species by refugia substrate were woody structures and leaf litter (Supplemental Table 6; Fig. 4). The positive slopes indicate that both were favored by Eastern American Toads. This model was evaluated for fit via residual analysis, and had a dispersion value of 1.07, with a p = 0.85. An examination of the QQ plot and residual vs predicted graph (Supplemental Figure) revealed some deviation, but this was still considered acceptable given the large sample size and low dispersion value.

**Figure 4.**
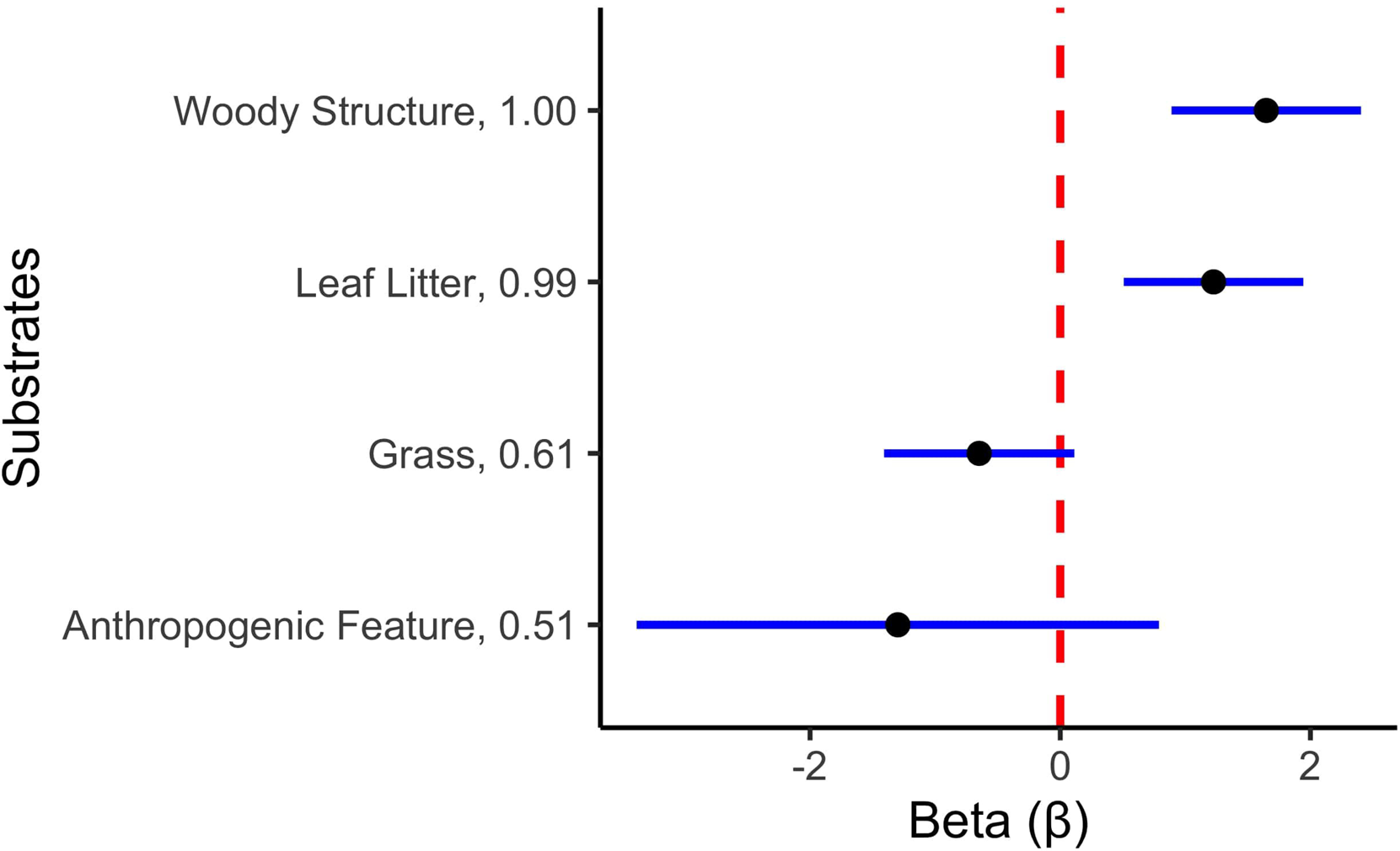
Model-averaged effect sizes and 95% confidence interval for all variables included in the final model of refugia substrate choice by toads. Variables are sorted from greatest to least AIC_c_ weight, which is reported after variable name. For the dependent variable of species, *Anaxyrus americanus americanus* was set to 1.

### Comparison of radiotelemetry and HDF

In comparing differences between initial moves and in the maximum detected move, whether a toad had a radiotelemetry tag was the strongest predictor variable. The model examining the number of days tracked between toads tagged with radiotelemetry or HDF did not perform any better than the null (Supplemental Tables 7-9).

## DISCUSSION

### Movement and tracking

Our findings indicate that for our species of interest, the recommended buffer of 30 meters in Virginia is insufficient for their movements (Virginia Department of Wildlife Resources, 2021). Both species’ median distance from breeding grounds was more than double this value. However, when considering long-distance movement, toads that were captured near or in a given forest patch stayed nearly entirely within it. In contrast, short-daily movement patterns included back-and-forth trips across forest edges. Less is known about juvenile emergence from their natal waters, and the pattern of directionality determination in their movements is still contentious (Patrick et al., 2007; Popescu et al., 2012; Walston & Mullin, 2008). As juveniles provide population connectivity via dispersal among amphibian populations, their survival and ability to travel prevents deleterious inbreeding effects (Semlitsch & Bodie, 2003).

In the movements of adult toads, our analyses indicated that high minimum temperatures were most important in driving larger movements by toads. This was similar to the results of other studies on amphibians living in environments with seasonal variability, due to their poikilothermic nature (Halstead et al., 2021; Henrique & Grant, 2019; Sinsch, 1988). Our results echo those of Sinsch (1988) in particular, which noted that temperatures approaching 0 °C correlated with decreasing movements in Common Toads (*Bufo bufo*). Three-day cumulative rainfall was also strongly related to movement, similar to results in movement and activity for other species of anuran (Fischer et al., 2020; Greenberg & Tanner, 2005). This is expected due to the constraining biology of toads as amphibians, as the permeability of their skin makes desiccation an activity restrictor. While toad skin is less susceptible to water loss than frog skin, the environmental availability of water is still a limiting factor for movement (Bentley & Yorio, 1976). Relative humidity was not retained in the final model. Some studies have found humidity to be linked to larger movements (D. J. Brown, 2013; Phillips et al., 2007), while others have found evidence that anurans select for refugia with high relative humidity (Z. L. Long & Prepas, 2012; Pitt et al., 2017). The capacity of anurans for independent selection of favorable local conditions, as well as the high average humidity in the Coastal Plain, likely explains the minimal importance of relative humidity in explaining overall activity levels and movement in our model. Overall, our results indicate that weather is an important driver of amphibian movement, with warm and rainy days providing the most favorable conditions for long-distance movements.

Landcover variables were also important in explaining the movements of both toad species. We found that toads moved shorter distances near trails, which we hypothesize may be because toads use these features for foraging. Previous work on the interactions between anurans and anthropogenic linear features, such as trails, has primarily focused on their ability to aid invasive toad dispersal or threaten connectivity (Brown et al., 2006; Cosentino et al., 2014). However, during captures, we noticed that toad densities were much higher on trails at night, and that parallel walks in adjacent woods yielded much fewer toad encounters. Combined with our results, we hypothesize that American and Fowler’s toads utilize trails as nocturnal hunting grounds due to their less complex substrate. Short movements are typically good indicators of foraging in many animal species, which aligns with our findings that toads move shorter distances near trails (Hooten et al., 2017). Furthermore, there is some support for this interaction in existing research, which found that Cane Toads (*Rhinella marina*) made more feeding attempts on insects on rough, lightly colored mats (González-Bernal et al., 2011), and all trails at our study sites (except for the paved bicycle trail at Greensprings Field) were made of a mixture of light-colored gravel and sand. Encounters on the paved bicycle trail could potentially be explained by toads using the asphalt trail at night for thermoregulation, or by differences in visual acuity between Cane Toads and our study species for hunting prey. In contrast, toads in areas with more coniferous landcover moved larger distances, suggesting a relative lack of habitat suitability in coniferous dominated areas compared to deciduous forests for our species of interest. Potential drivers behind these greater movements may include higher soil acidity creating irritating conditions for toads (Wyman, 1990), or a lack of sufficient vegetative refugia, as the allelopathic nature of pine needles can suppress undergrowth (Duryea et al., 1999; Mallik, 1998). Both variables are avenues for future inquiry that could be manipulated in future studies to further identify toad preferences between substrate types and available landcover features, and the exact mechanisms behind them.

We found that radio telemetry allows for toads to be tracked over longer distances, though the days an individual is tracked did not differ between radiotelemetry and HDF tag types. Radiotelemetry tags were associated with longer initial distances, though this may be because toads with radiotelemetry tags are more reliably tracked over greater distances, while HDF tagged toads are lost. In the 2023 field season when we used radiotelemetry, 9/9 lost toads had HDF tags. Alternatively, it may be due to greater stress associated with equipping toads with heavier tags. An endocrinal analysis of stress levels in tagged toads could help differentiate between these hypotheses (Bliley & Woodley, 2012). Our findings of differences between tag types echo those of Alford & Rowley (2007), who found that radiotagged toads were relocated more frequently and at greater distances, further encouraging scientists to consider the cost and effort tradeoffs between HDF and radiotelemetry tracking methods. If others wish to combine these methods as well, a metal detector could achieve a similar result to the RECCO detector, and for a lower price.

### Site fidelity

Our study represents one of the first to employ random walks to evaluate site fidelity to their refugia during the nonbreeding season. Site fidelity has been broadly observed in amphibians during studies of life history, but our results provide quantitative evidence to those observations during the adult non-breeding stage (Gamble et al., 2007; Luger et al., 2009; Pittman et al., 2008). These findings emphasize that in addition to breeding site fidelity, at least some species of anurans may also be characterized by refugia fidelity. The theory of site fidelity posits that staying in a habitat with a known suite of refugia sites and resources is conducive to survival, leading to greater fitness than movements that increase predation risk and waste energy (Sanz-Aguilar et al., 2012). These assumptions play out for our study species, but at a smaller spatial scale than the model systems of birds in which this theory has mainly been described. Unlike birds, established adult amphibians do not establish genetic connectivity by large movements. Instead, amphibian genetic connectivity is established by juvenile dispersal post-metamorphosis (Pittman et al., 2014; Semlitsch, 2008). These differences explain annual amphibian migratory behavior, as well as daily returns to previously occupied refugia.

### Substrate use

We found differences between the two species in their substrate choice for daytime refuge. Notably, Eastern American Toads favored natural wood structures and leaf litter for shelter, while Fowler’s Toads appeared to have no strong preferences for any substrate type. Coarse woody debris and leaf litter are a commonly described refugia type for other species of toads, due to the combination of climatic insulation and predator protection that they provide (Browne & Paszkowski, 2018; Lemckert & Slatyer, 2002; Long & Prepas, 2012; Pitt et al., 2013; Seebacher & Alford, 2002).

While not associated with any one species in the final model, anthropogenic structure use was encountered at a relatively steady rate throughout the study. The main structures that were utilized were foot bridges and drainage pipes that were originally placed to prevent path wash-out. These structures were used nearly entirely by Fowler’s Toads (73/74 observations), though this likely reflects site availability, as these features were most common at Greensprings Trail, where Fowler’s Toads were most abundant. Additionally, communal use of refugia was also observed in Fowler’s Toads, as tagged toads were sometimes found to be cohabitating in log cavities with 1-2 other toads.

Our study found that Fowler’s Toads are more of a habitat generalist compared to Eastern American Toads. This finding is unusual in the context of reports of steady declines in populations of Fowler’s Toads, as it is generally expected that more specialized species are at a greater risk for population decline (Clavel et al., 2011; Nowakowski et al., 2018). The most likely current explanation may be the one that Jones & Tupper, (2015) suggest, in that Fowler’s Toads habitat selection does not occur at the microhabitat level, but instead the overall suitability of the macrohabitat. Their analysis suggested that Fowler’s Toads rely on sparse forest cover and low-intensity agricultural activity in the surrounding area. Because the current forest patches are maturing successionally and agricultural pressure is intensifying due to transitions away from local farming, these changes may explain the long-term trend of decline (McGarvey et al., 2015; Vogeler, 2019). Further investigations to identify and compare areas of maintenance to areas of decline for Fowler’s Toads populations are needed, to better elucidate the differences between these otherwise similar species of toads.

### Management implications

Our results suggest that both micro- and macrohabitat are of crucial importance to anuran species. Both types of habitats must be maintained by land managers who wish to maintain healthy populations, such as by leaving woody debris and leaf litter on the forest floor, and by ensuring appropriate habitat types for species of concern that are present. The activity of toads on paths also suggests that managers should take care to warn any nocturnal visitors to their grounds that they should watch their step. While “Big Night” amphibian migration events do not generally occur in this region, our research and personal observations indicates that paths within protected areas are a major attractant of toads throughout their active period of the year (Hedrick et al., 2019). Our findings also have broader implications in the context of climate change, as the importance of environmental variables indicates the importance of climate and the weather it produces on amphibian behavior. Our own results are opposed in this context though, as while minimum temperatures are expected to increase, rainfall is predicted to become increasingly sporadic and intense (Lee et al., 2023). However, when placed within the context of the larger body of research on amphibian sensitivity to desiccation and our own findings in relation to how cumulative 3-day rainfall relates to movement, we agree with other researchers that climate change will likely lead to decreased activity, and potential extirpation of populations if the risks of foraging activity away from refugia are increased above the benefits (Blaustein et al., 2010; Lertzman-Lepofsky et al., 2020). To increase resilience of amphibians to the current threats leveled at them, we encourage a prioritization of protection for large and connected areas for amphibians of concern, of both their breeding and upland habitats at scales that are biologically relevant (Kremen & Merenlender, 2018; Timmers et al., 2022).

## DATA ACCESSIBILITY

Supplemental material is available at https://www.ichthyologyandherpetology.org/XXX.

## Acknowledgments

We would like to thank Millan Khadka, Chris Greene, Jill Ashey, Simeon Brown, Joseph Moriarty, Jesse Smyth, Trent Stafford, Matt Whalen, Madison White, Alexa Busby, Addy Cullen, and Lauren Scherrer for their assistance in capturing and tagging toads at night. We would also like to thank the Martin family for giving us access to their property, Tom Meyer for technical advice on belt construction, Drew LaMar for troubleshooting R code, and Jennifer Swenson for supplying us with NASA Planet data. Funding was provided by the William & Mary Department of Biology’s Ferguson Undergraduate Research Fund, Plumeri Award, and the Charles Center.

---

Supplemental Table 1: Crosswalk of Virginia Land Cover (LC) Dataset

Supplemental Table 2: Crosswalk of Substrate categories. Numbers in parentheses present the number of observations in each substrate

Supplemental Table 3: Evaluation of candidate variables on the basis of the information theoretic approach. Variables with an asterisk were carried forward to the final model. If top variables with multiple threshold values were within 2 AIC of each other, the one with the lowest AIC score was carried forward to the final model. Threshold column indicates the different levels at which a variable was evaluated (1^st^, median, and 3rd indicates the level was at the 1^st^, median, and 3rd quartile of non-zero movements; NA means there was no threshold level). Extent/Decay column indicates whether a given variable was examined at three possible extent levels (E), or at four levels of variable forms: three levels of decay (D) and the linear form (L). For each model, we show degrees of freedom (df), AIC values adjusted for small samples sizes (AIC**_c_**), and the delta AIC values (Δ AIC_c_) in comparison to the null model (AICc = 6538.005).

Supplemental Table 4: Effect sizes for six final candidate predictor variables estimated in all possible model combinations (n = 126, not including null model). The null model (second to last row) and Julian day model (on last row, with Δ AIC = 1.8) were not included in model-averaged effect-size estimates for the final model. A “*NA*” indicates that a given predictor variable was not included in a given model. For each model, we show degrees of freedom (df), log likelihood estimates (logLik), AIC values adjusted for small samples sizes (AIC**_c_**), delta AIC values (Δ AIC_c_), and the AIC weights (**ω**).

Supplemental Table 5: Evaluation of candidate predictor variables on the basis of the information-theoretic approach. Models with an asterisk were included in the final model on the basis of being 2 AIC units or less compared to the null. For each model, we show degrees of freedom (df), AIC values adjusted for small samples sizes (AIC**_c_**), and the delta AIC values (Δ AIC_c_) in comparison to the null (AICc = 858.32).

Supplemental Table 6: Effect sizes for 4 final candidate predictor variables estimated in all possible model combinations (n = 16, not including null model). The null model (bottom row) was not included in model-averaged effect-size estimates for the final model. A “*NA*” indicates that a given predictor variable was not included in a given model. For each model, we show degrees of freedom (df), log likelihood estimates (logLik), AIC values adjusted for small samples sizes (AIC**_c_**), delta AIC values (Δ AIC_c_), and the AIC weights (**ω**).

Supplemental Figure 1: QQ plot of residual and residual vs. predicted plot for the final movement analysis model. Despite the report of significant deviations, visual inspection reveals a close fit to the expected line in the QQ plot, and no sign of funneling in the residual vs. predicted graph, allowing us to accept the final model.

Supplemental Figure 2: QQ plot of residual and residual vs. predicted plot for the final substrate analysis model. Some deviation tests reported as significant, but the deviation still appears to be acceptable visually for our model choice.

Supplemental Table 7: Model selection for variables relating to the distance travelled after initial tagging for HDF and radiotelemetry. Radiotelemetry tagged toads were set to 1.

Supplemental Table 8: Model selection for variables relating the maximum distance toads were found between relocations.

Supplemental Table 9: Model selection for variables relating the number of days toads were tracked for

