## Supplemental Table 1 for "A Combination of Environmental and Landscape Variables Drive Movement and Habitat use in Two *Anaxyrus* sp. in the Eastern Coastal Plain"

| **Virginia LC name** | **Original value** | **New values** | **New name** |
| --- | --- | --- | --- |
| Conifer Forest | 1 | 1 | Conifer Forest |
| Open Water | 11 | 11 | Open Water |
| Extracted Impervious | 21 | 21 | Impervious |
| External Impervious | 22 | 21 | Impervious |
| Barren | 31 | 31 | Barren |
| Forest | 41 | 42 | Forest |
| Tree | 42 | 42 | Forest |
| Scrub/Shrub | 51 | 51 | Scrub/Shrub |
| Harvested/Disturbed | 61 | 61 | Harvested/Disturbed |
| Turf/Grass | 71 | 71 | Grass |
| Pasture | 81 | 71 | Grass |
| Cropland | 82 | 82 | Cropland |
| NWI | 91 | 91 | NWI |
