## Supplemental Table 2 for "A Combination of Environmental and Landscape Variables Drive Movement and Habitat use in Two *Anaxyrus* sp. in the Eastern Coastal Plain"

| **Name (n)** | **Original substrate code** | **Reclassed substrate code** | **Reclassed substrate name** |
| --- | --- | --- | --- |
| Bare ground (9) | bg | b | bare ground |
| Bare sandy ground (2) | bgs | b | bare ground |
| Tree cavity (102) | ca | cwd | Tree feature |
| Coarse Woody Debris, in cover (168) | cwdc | cwd | Tree feature |
| Coarse Woody Debris, on-top (7) | cwdt | cwd | Tree feature |
| Drainpipe (36) | dp | anth | Anthropogenic feature |
| Drain spout (1) | dsb | anth | Anthropogenic feature |
| Dried aquatic vegetation (2) | gav | g | grass |
| Mixed grass, stiltgrass and non-stiltgrass (6) | gm | g | grass |
| Grass non-stiltgrass (331) | gn | g | grass |
| Half buried, with grassy cover (3) | gnsb | g | grass |
| Grass, stiltgrass (19) | gs | g | grass |
| Leaf litter buried (497) | lfb | L | Leaf Litter |
| Mixed leaf litter buried (22) | lfbpn | L | Leaf Litter |
| On top of leaf litter (26) | lft | L | Leaf Litter |
| On top of pine leaf litter (1) | lftpn | L | Leaf Litter |
| Moss (40) | m | m | Moss |
| On top of pine needles (2) | pn | L | Leaf Litter |
| Half buried in pine needles (3) | pnsb | L | Leaf Litter |
| buried in sand (14) | s | s | Buried in substrate |
| soil buried (158) | sb | s | Buried in substrate |
| Small mammal burrow (6) | smb | s | Buried in substrate |
| Tree trunk (2) | tt | cwd | Tree feature |
| Under foot bridge (37) | ubr | anth | Anthropogenic feature |
| Water (3) | w | w | Water |
