## Supplemental Table 3 for "A Combination of Environmental and Landscape Variables Drive Movement and Habitat use in Two *Anaxyrus* sp. in the Eastern Coastal Plain"

| **Model** | **Threshold** | **Extent/Decay** | **df** | **AICc** | **ΔAIC** |
| --- | --- | --- | --- | --- | --- |
| Cumulative rainfall | 3 days* | NA | 4 | 6531.39 | 0.00 |
|  | 4 days | NA | 4 | 6531.78 | 0.39 |
|  | 5 days | NA | 4 | 6532.03 | 0.64 |
|  | 2 days | NA | 4 | 6532.35 | 0.96 |
|  | 1 day | NA | 4 | 6533.08 | 1.69 |
|  | 6 days | NA | 4 | 6534.86 | 3.47 |
|  | 7 days | NA | 4 | 6536.86 | 5.47 |
|  | 14 days | NA | 4 | 6537.38 | 5.99 |
| Daily temperature | Minimum* | NA | 4 | 6527.23 | 0 |
|  | Mean | NA | 4 | 6534.38 | 7.15 |
|  | Maximum | NA | 4 | 6539.02 | 11.79 |
| Average relative humidity | NA | NA | 4 | 6538.4 | 0.39 |
| Proportion of Forest | Median | E | 4 | 6538.08 | 0.07 |
|  | 1st | E | 4 | 6538.21 | 0.20 |
|  | 3rd | E | 4 | 6538.32 | 0.32 |
| Proportion conifer | 3rd* | E | 4 | 6533.56 | 0.00 |
|  | Median | E | 4 | 6536.14 | 2.58 |
|  | 1st | E | 4 | 6537.22 | 3.66 |
| TRI | 1st | E | 4 | 6537.18 | 0.00 |
|  | Median | E | 4 | 6537.48 | 0.30 |
|  | 3rd | E | 4 | 6539.16 | 1.98 |
| Distance to water | NA* | L | 4 | 6536.32 | 0.00 |
|  | Median | D | 4 | 6538.22 | 1.90 |
|  | 1st | D | 4 | 6538.52 | 2.20 |
|  | 3rd | D | 4 | 6539.43 | 3.11 |
| Distance to trail | NA* | L | 4 | 6528.69 | 0.00 |
|  | 1st | D | 4 | 6538.23 | 9.54 |
|  | 3rd | D | 4 | 6538.64 | 9.95 |
|  | Median | D | 4 | 6539.87 | 11.18 |
| Distance to breeding site | NA | L | 4 | 6538.57 | 0.57 |
|  | 3rd | D | 4 | 6539.79 | 1.78 |
|  | Median | D | 4 | 6539.79 | 1.79 |
|  | 1st | D | 4 | 6539.81 | 1.81 |
| Radiotagged | NA | NA | 4 | 6539.79 | 1.78 |
