## Supplemental Table 4 for "A Combination of Environmental and Landscape Variables Drive Movement and Habitat use in Two *Anaxyrus* sp. in the Eastern Coastal Plain"

| **Intercept** | **Julian Day** | **% Conifer 3Q** | **3-day Rain** | **Distance to trail** | **Species** | **Minimum Daily Temperature** | **df** | **logLik** | **AICc** | **delta** | **weight** |
| --- | --- | --- | --- | --- | --- | --- | --- | --- | --- | --- | --- |
| 1.65 | -0.12 | 0.74 | 0.17 | 0.18 | -0.28 | 0.32 | 10.00 | -3243.02 | 6506.19 | 0.00 | 3.53E-01 |
| 1.66 | *NA* | 0.77 | 0.16 | 0.18 | -0.30 | 0.25 | 9.00 | -3244.51 | 6507.15 | 0.96 | 2.18E-01 |
| 1.52 | -0.14 | 0.96 | 0.17 | 0.17 | *NA* | 0.33 | 9.00 | -3244.88 | 6507.88 | 1.69 | 1.52E-01 |
| 1.78 | -0.13 | *NA* | 0.18 | 0.19 | -0.40 | 0.32 | 9.00 | -3245.26 | 6508.64 | 2.44 | 1.04E-01 |
| 1.52 | *NA* | 0.98 | 0.16 | 0.17 | *NA* | 0.25 | 8.00 | -3246.69 | 6509.48 | 3.28 | 6.83E-02 |
| 1.80 | *NA* | *NA* | 0.16 | 0.19 | -0.43 | 0.24 | 8.00 | -3246.95 | 6509.99 | 3.80 | 5.28E-02 |
| 1.61 | -0.14 | *NA* | 0.17 | 0.18 | *NA* | 0.31 | 8.00 | -3248.70 | 6513.49 | 7.29 | 9.19E-03 |
| 1.67 | *NA* | 0.78 | *NA* | 0.18 | -0.29 | 0.27 | 8.00 | -3248.90 | 6513.90 | 7.71 | 7.48E-03 |
| 1.66 | -0.09 | 0.75 | *NA* | 0.18 | -0.27 | 0.32 | 9.00 | -3248.17 | 6514.45 | 8.26 | 5.67E-03 |
| 1.67 | -0.11 | 0.74 | 0.17 | *NA* | -0.28 | 0.34 | 9.00 | -3248.45 | 6515.03 | 8.84 | 4.25E-03 |
| 1.61 | *NA* | *NA* | 0.16 | 0.18 | *NA* | 0.23 | 7.00 | -3250.56 | 6515.19 | 9.00 | 3.92E-03 |
| 1.68 | *NA* | 0.77 | 0.15 | *NA* | -0.30 | 0.28 | 8.00 | -3249.74 | 6515.57 | 9.38 | 3.24E-03 |
| 1.55 | -0.12 | 1.00 | *NA* | 0.18 | *NA* | 0.32 | 8.00 | -3249.76 | 6515.62 | 9.42 | 3.17E-03 |
| 1.54 | *NA* | 0.99 | *NA* | 0.17 | *NA* | 0.26 | 7.00 | -3250.84 | 6515.75 | 9.56 | 2.97E-03 |
| 1.54 | -0.13 | 0.95 | 0.16 | *NA* | *NA* | 0.34 | 8.00 | -3250.25 | 6516.60 | 10.41 | 1.94E-03 |
| 1.81 | *NA* | *NA* | *NA* | 0.19 | -0.42 | 0.26 | 7.00 | -3251.39 | 6516.85 | 10.66 | 1.71E-03 |
| 1.80 | -0.10 | *NA* | *NA* | 0.18 | -0.39 | 0.32 | 8.00 | -3250.48 | 6517.06 | 10.87 | 1.54E-03 |
| 1.70 | *NA* | 0.70 | 0.17 | 0.20 | -0.28 | *NA* | 8.00 | -3250.71 | 6517.52 | 11.33 | 1.22E-03 |
| 1.81 | -0.12 | *NA* | 0.17 | *NA* | -0.40 | 0.34 | 8.00 | -3250.79 | 6517.67 | 11.48 | 1.13E-03 |
| 1.54 | *NA* | 0.98 | 0.15 | *NA* | *NA* | 0.27 | 7.00 | -3251.85 | 6517.77 | 11.58 | 1.08E-03 |
| 1.60 | *NA* | 0.91 | 0.16 | 0.19 | *NA* | *NA* | 7.00 | -3252.13 | 6518.33 | 12.14 | 8.15E-04 |
| 1.82 | *NA* | *NA* | 0.16 | *NA* | -0.43 | 0.26 | 7.00 | -3252.29 | 6518.66 | 12.46 | 6.93E-04 |
| 1.81 | *NA* | *NA* | 0.17 | 0.20 | -0.39 | *NA* | 7.00 | -3252.72 | 6519.51 | 13.32 | 4.52E-04 |
| 1.70 | 0.01 | 0.70 | 0.17 | 0.20 | -0.28 | *NA* | 9.00 | -3250.71 | 6519.54 | 13.34 | 4.46E-04 |
| 1.60 | -0.03 | 0.92 | 0.16 | 0.20 | *NA* | *NA* | 8.00 | -3252.07 | 6520.23 | 14.04 | 3.15E-04 |
| 1.81 | 0.01 | *NA* | 0.17 | 0.20 | -0.39 | *NA* | 8.00 | -3252.71 | 6521.51 | 15.32 | 1.66E-04 |
| 1.64 | -0.11 | *NA* | *NA* | 0.18 | *NA* | 0.31 | 7.00 | -3253.77 | 6521.62 | 15.42 | 1.58E-04 |
| 1.69 | *NA* | 0.77 | *NA* | *NA* | -0.28 | 0.30 | 7.00 | -3253.84 | 6521.76 | 15.56 | 1.47E-04 |
| 1.64 | *NA* | *NA* | *NA* | 0.18 | *NA* | 0.25 | 6.00 | -3254.86 | 6521.77 | 15.58 | 1.46E-04 |
| 1.64 | -0.13 | *NA* | 0.17 | *NA* | *NA* | 0.33 | 7.00 | -3254.22 | 6522.52 | 16.32 | 1.01E-04 |
| 1.68 | -0.08 | 0.75 | *NA* | *NA* | -0.27 | 0.34 | 8.00 | -3253.21 | 6522.52 | 16.33 | 1.00E-04 |
| 1.67 | *NA* | *NA* | 0.17 | 0.20 | *NA* | *NA* | 6.00 | -3255.58 | 6523.22 | 17.03 | 7.07E-05 |
| 1.56 | *NA* | 0.98 | *NA* | *NA* | *NA* | 0.29 | 6.00 | -3255.69 | 6523.43 | 17.24 | 6.36E-05 |
| 1.56 | -0.10 | 0.96 | *NA* | *NA* | *NA* | 0.34 | 7.00 | -3254.79 | 6523.66 | 17.46 | 5.69E-05 |
| 1.64 | *NA* | *NA* | 0.15 | *NA* | *NA* | 0.25 | 6.00 | -3255.91 | 6523.88 | 17.69 | 5.09E-05 |
| 1.76 | *NA* | 0.85 | *NA* | 0.20 | -0.28 | *NA* | 7.00 | -3255.22 | 6524.51 | 18.32 | 3.71E-05 |
| 1.63 | *NA* | 0.97 | *NA* | 0.19 | *NA* | *NA* | 6.00 | -3256.37 | 6524.79 | 18.60 | 3.23E-05 |
| 1.83 | *NA* | *NA* | *NA* | *NA* | -0.41 | 0.28 | 6.00 | -3256.43 | 6524.91 | 18.72 | 3.03E-05 |
| 1.68 | -0.02 | *NA* | 0.17 | 0.20 | *NA* | *NA* | 7.00 | -3255.55 | 6525.18 | 18.99 | 2.66E-05 |
| 1.82 | -0.09 | *NA* | *NA* | *NA* | -0.39 | 0.34 | 7.00 | -3255.62 | 6525.31 | 19.12 | 2.48E-05 |
| 1.76 | 0.01 | 0.84 | *NA* | 0.20 | -0.28 | *NA* | 8.00 | -3255.22 | 6526.53 | 20.34 | 1.35E-05 |
| 1.63 | 0.00 | 0.97 | *NA* | 0.19 | *NA* | *NA* | 7.00 | -3256.37 | 6526.81 | 20.61 | 1.18E-05 |
| 1.85 | *NA* | *NA* | *NA* | 0.20 | -0.39 | *NA* | 6.00 | -3257.72 | 6527.49 | 21.30 | 8.36E-06 |
| 1.71 | *NA* | 0.65 | 0.16 | *NA* | -0.28 | *NA* | 7.00 | -3257.21 | 6528.49 | 22.29 | 5.08E-06 |
| 1.84 | 0.04 | *NA* | *NA* | 0.20 | -0.39 | *NA* | 7.00 | -3257.59 | 6529.26 | 23.07 | 3.45E-06 |
| 1.62 | *NA* | 0.89 | 0.16 | *NA* | *NA* | *NA* | 6.00 | -3258.66 | 6529.37 | 23.18 | 3.27E-06 |
| 1.66 | *NA* | *NA* | *NA* | *NA* | *NA* | 0.27 | 5.00 | -3259.87 | 6529.78 | 23.58 | 2.67E-06 |
| 1.66 | -0.10 | *NA* | *NA* | *NA* | *NA* | 0.33 | 6.00 | -3258.87 | 6529.79 | 23.59 | 2.65E-06 |
| 1.71 | 0.04 | 0.67 | 0.16 | *NA* | -0.28 | *NA* | 8.00 | -3257.05 | 6530.20 | 24.01 | 2.16E-06 |
| 1.83 | *NA* | *NA* | 0.17 | *NA* | -0.38 | *NA* | 6.00 | -3259.11 | 6530.27 | 24.08 | 2.09E-06 |
| 1.71 | *NA* | *NA* | *NA* | 0.20 | *NA* | *NA* | 5.00 | -3260.19 | 6530.42 | 24.23 | 1.93E-06 |
| 1.62 | 0.01 | 0.88 | 0.16 | *NA* | *NA* | *NA* | 7.00 | -3258.65 | 6531.38 | 25.19 | 1.20E-06 |
| 1.83 | 0.03 | *NA* | 0.16 | *NA* | -0.39 | *NA* | 7.00 | -3259.03 | 6532.13 | 25.94 | 8.23E-07 |
| 1.71 | 0.01 | *NA* | *NA* | 0.20 | *NA* | *NA* | 6.00 | -3260.19 | 6532.43 | 26.24 | 7.08E-07 |
| 1.69 | *NA* | *NA* | 0.16 | *NA* | *NA* | *NA* | 5.00 | -3262.01 | 6534.06 | 27.86 | 3.14E-07 |
| 1.80 | *NA* | 0.84 | *NA* | *NA* | -0.30 | *NA* | 6.00 | -3261.37 | 6534.80 | 28.61 | 2.16E-07 |
| 1.67 | *NA* | 0.94 | *NA* | *NA* | *NA* | *NA* | 5.00 | -3262.60 | 6535.25 | 29.05 | 1.73E-07 |
| 1.68 | 0.02 | *NA* | 0.16 | *NA* | *NA* | *NA* | 6.00 | -3261.99 | 6536.03 | 29.84 | 1.17E-07 |
| 1.72 | 0.07 | 0.69 | *NA* | *NA* | -0.27 | *NA* | 7.00 | -3261.41 | 6536.88 | 30.69 | 7.63E-08 |
| 1.66 | 0.02 | 0.94 | *NA* | *NA* | *NA* | *NA* | 6.00 | -3262.55 | 6537.16 | 30.97 | 6.64E-08 |
| 1.85 | *NA* | *NA* | *NA* | *NA* | -0.37 | *NA* | 5.00 | -3263.85 | 6537.75 | 31.56 | 4.95E-08 |
| 1.85 | 0.06 | *NA* | *NA* | *NA* | -0.38 | *NA* | 6.00 | -3263.49 | 6539.04 | 32.85 | 2.60E-08 |
| 1.74 | *NA* | *NA* | *NA* | *NA* | *NA* | *NA* | 4.00 | -3266.35 | 6540.72 | 34.53 | 1.12E-08 |
| 1.72 | 0.03 | *NA* | *NA* | *NA* | *NA* | *NA* | 5.00 | -3266.24 | 6542.52 | 36.33 | 4.56E-09 |
