## Supplemental Table 5 for "A Combination of Environmental and Landscape Variables Drive Movement and Habitat use in Two *Anaxyrus* sp. in the Eastern Coastal Plain"

| **Model** | **df** | **AIC_c_** | **ΔAIC_c_** |
| --- | --- | --- | --- |
| Anthropogenic | 3 | 851.58 | 0.00 |
| Bare | 3 | 857.65 | 0.00 |
| Grass | 3 | 828.29 | 0.00 |
| Leaf | 3 | 848.17 | 0.00 |
| Moss | 3 | 860.09 | 1.77 |
| Soil | 3 | 858.82 | 0.50 |
| Water | 3 | 859.82 | 1.50 |
| Wood | 3 | 836.77 | 0.00 |
