## Supplemental Table 6 for "A Combination of Environmental and Landscape Variables Drive Movement and Habitat use in Two *Anaxyrus* sp. in the Eastern Coastal Plain"

| **Intercept** | **Anthropogenic** | **Grass** | **Leaf** | **Woody** | **df** | **logLik** | **AIC_c_** | **ΔAIC_c_** | **ω** |
| --- | --- | --- | --- | --- | --- | --- | --- | --- | --- |
| 0.77 | -1.41 | -0.71 | 0.99 | 1.42 | 6 | -395.67 | 803.39 | 0.00 | 0.34 |
| 0.60 | *NA* | -0.54 | 1.19 | 1.63 | 5 | -396.98 | 804.00 | 0.61 | 0.25 |
| 0.29 | *NA* | *NA* | 1.48 | 1.91 | 4 | -398.08 | 804.18 | 0.79 | 0.23 |
| 0.34 | -1.01 | *NA* | 1.41 | 1.83 | 5 | -397.44 | 804.92 | 1.53 | 0.16 |
| 1.45 | -2.10 | -1.34 | *NA* | 0.73 | 5 | -400.47 | 810.97 | 7.58 | 0.01 |
| 1.37 | *NA* | -1.26 | *NA* | 0.86 | 4 | -404.67 | 817.36 | 13.97 | 0.00 |
| 1.62 | -2.36 | -1.50 | *NA* | *NA* | 4 | -405.08 | 818.18 | 14.79 | 0.00 |
| 1.56 | -2.31 | -1.45 | 0.12 | *NA* | 5 | -404.95 | 819.93 | 16.54 | 0.00 |
| 1.57 | *NA* | -1.44 | *NA* | *NA* | 3 | -411.15 | 828.31 | 24.92 | 0.00 |
| 1.43 | *NA* | -1.32 | 0.27 | *NA* | 4 | -410.43 | 828.89 | 25.50 | 0.00 |
| 0.98 | -1.77 | *NA* | *NA* | 1.06 | 4 | -412.70 | 833.44 | 30.05 | 0.00 |
| 0.95 | *NA* | *NA* | *NA* | 1.15 | 3 | -415.39 | 836.79 | 33.40 | 0.00 |
| 0.92 | -1.87 | *NA* | 0.64 | *NA* | 4 | -417.95 | 843.93 | 40.54 | 0.00 |
| 0.87 | *NA* | *NA* | 0.72 | *NA* | 3 | -421.08 | 848.19 | 44.80 | 0.00 |
| 1.15 | -2.10 | *NA* | *NA* | *NA* | 3 | -422.79 | 851.59 | 48.21 | 0.00 |
| 1.12 | *NA* | *NA* | *NA* | *NA* | 2 | -427.16 | 858.33 | 54.94 | 0.00 |
