## Supplementary figures and images for "A Combination of Environmental and Landscape Variables Drive Movement and Habitat use in Two *Anaxyrus* sp. in the Eastern Coastal Plain"

### Supplemental Figure 1

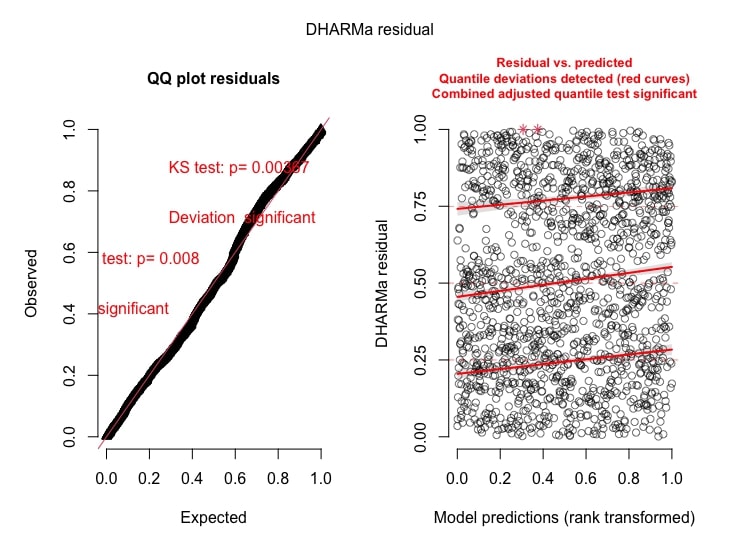

### Supplemental Figure 2

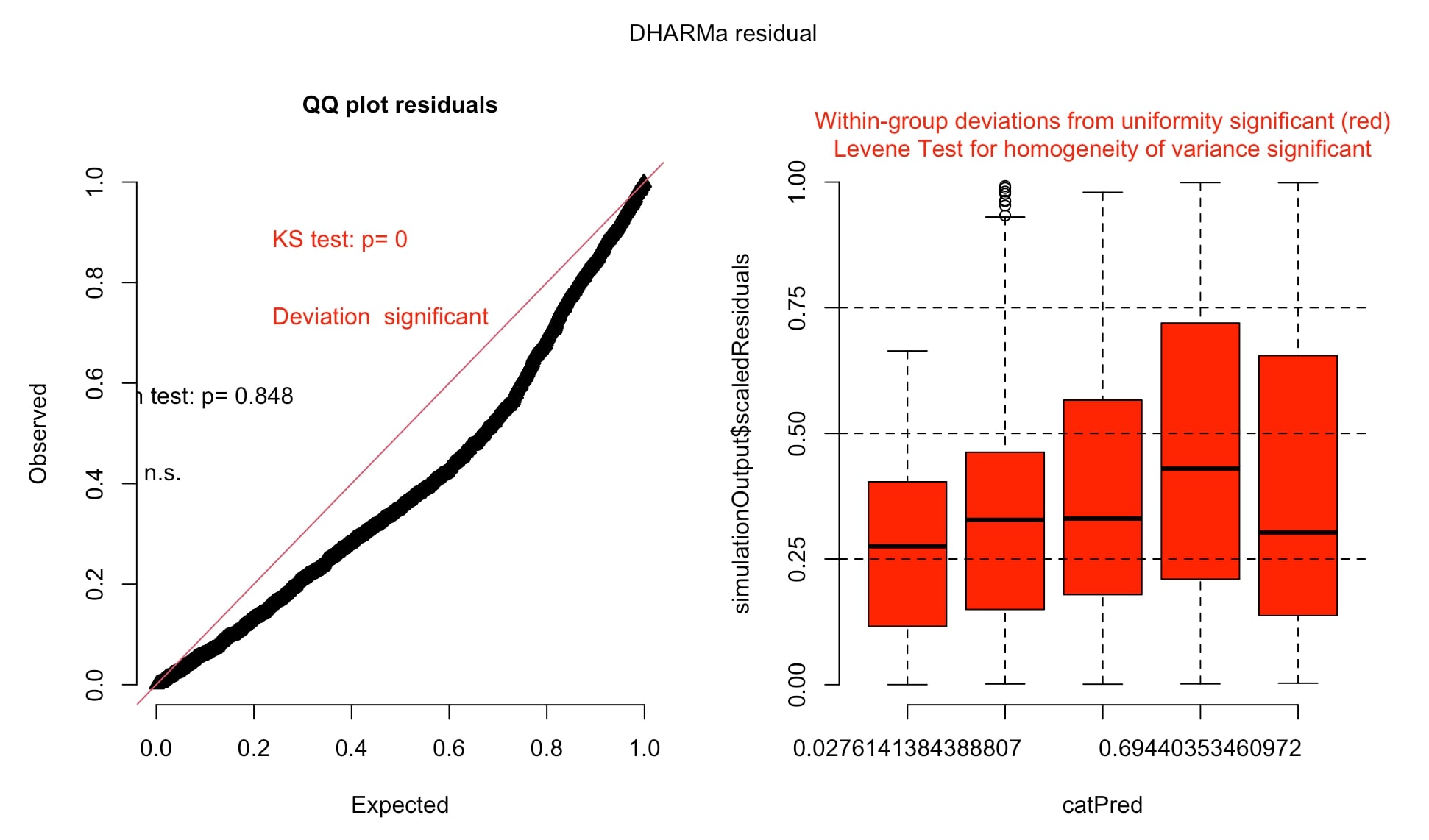
